# Mapping function in the tree shrew visual system using functional ultrasound imaging

**DOI:** 10.1101/2025.10.03.680247

**Authors:** Joseph B. Wekselblatt, Rohit Nayak, Frank F. Lanfranchi, Francisco Luongo

## Abstract

We adapted functional ultrasound imaging (fUSI) for awake, head-fixed northern tree shrews and used it to produce functional maps of visual processing at ∼100 µm spatial resolution and ∼100 ms temporal resolution. Using classical retinotopic stimuli, full-field noise, motion localizers, and object versus scrambled object contrasts, we demonstrate robust, spatially specific hemodynamic responses across primary and extrastriate visual cortex, superior colliculus and subcortical structures. fUSI reliably reveals retinotopic reversals, laterality, and stimulus-selective modules, and yields high signal-to-noise %CBV changes that enable single-session mapping and targeting of electrophysiology or perturbations. These mesoscale maps provide a systems-level complement to recent high-density electrophysiological surveys of tree shrew visual cortex, which reported a compressed ventral-stream hierarchy and surprisingly early emergence of object coding in V2 (Lanfranchi et al., 2025). Together, our results establish fUSI as a powerful, scalable tool for functional mapping in the tree shrew, bridging large-scale circuit measurement and single-neuron electrophysiology and accelerating this species’ utility as a bridge between rodent genetics and primate vision.

## Introduction

The choice and assessment of model species is an important consideration in biomedical studies, particularly when such models are intended to generate knowledge that will translate to humans. The challenge of finding adequate non-human models for translational research is particularly acute in neuroscience: neurobiological and behavioral phenotypes are complex and plastic, and many traits important in humans are absent, radically different, or difficult to assess in other animals. The most common mammalian species for mechanistic neuroscience research fall into two main categories: non-human primates and rodent species. Existing animal models often present methodological challenges: in primates, which allow the probing of complex cognition, it is difficult to record from large ensembles of neurons across multiple brain areas or manipulate genetically defined cell populations; in rodents, rudimentary cortical organization and behavioral repertoire limits the modeling of visual processing, and the acuity of rodent vision is ∼2-3 orders of magnitude worse than humans.

For these reasons, we chose to study visual organization in the tree shrew, an animal with high visual acuity and considerable cognitive abilities. The tree shrew is closely related to primates (Lee, Huang, & Fitzpatrick, 2016; Xiao, Liu, & Chen, 2017). A diurnal animal, it has high visual acuity (>10x that of rodents) and strong visual form discrimination abilities (Petry, Fox, & Casagrande, 1984; Schumacher, McCann, Maximov, & Fitzpatrick, 2021). Neural hardware-wise, it has a cone-dominant retina (>95% cones), a visual cortex organized into functional columns including orientation pinwheels, and a large fraction of cortex devoted to visual form processing (Lyon, Jain, & Kaas, 1998; Petry & Bickford, 2019; Sajdak et al., 2019; Wong & Kaas, 2009). Moreover, unlike the macaque, tree shrew cortex is lissencephalic (no sulci or gyri), allowing increased access for imaging and electrophysiological recording of neural circuit dynamics. Due to its small size and quick developmental and reproductive cycles (4-6 months from birth to adulthood, with 2-4 offspring born each litter every 2 months), it offers experimental and genetic accessibility similar to rodents. The tree shrew has been used widely as a model for cone disorders, myopia, and other diseases of the eye (Timothy J. Gawne, Siegwart, Ward, & Norton, 2017; T. J. Gawne, Ward, & Norton, 2018; Sajdak et al., 2019; Ward, Norton, Huisingh, & Gawne, 2018). Tree shrews have similar visual acuity to cats and ferrets, with a smaller body size than rats, making them an ideal organism for mechanistic studies in visual neuroscience (Jang, Song, & Paik, 2019).

To explore the visual cortical organization of this highly visual, and under-utilized species we employed functional ultrasound imaging. Functional ultrasound imaging (fUSI) is an emerging technology for mapping cerebral blood volume changes, related to brain activity through neurovascular coupling, <u>across the entire brain with high spatiotemporal resolution (100 μm, ∼100 ms) (Figure 1)</u>. To date, fUSI has been used to image activity of 3-D brain volumes in mice (Mace et al., 2018; C. Rabut, Ferrier, et al., 2020; Tiran et al., 2017), rats (Claire Rabut et al., 2019; Sieu et al., 2015; Urban et al., 2014), ferrets (Demené et al., 2016; Hu, Zhu, Briggs, & Doyley, 2023), non-human primates (Norman et al., 2020), and human patients (Demene et al., 2017; Imbault, Chauvet, Gennisson, Capelle, & Tanter, 2017; Topcuoglu, Aydin, & Saka, 2009).

**Figure 1).**
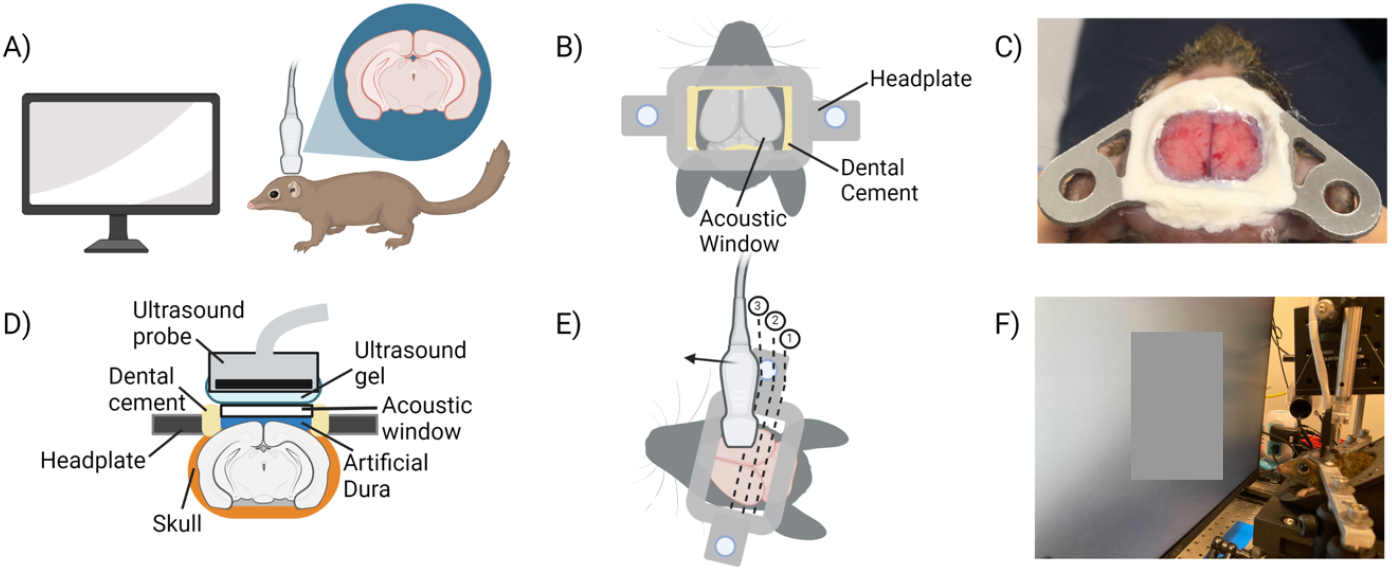
Chronic acoustic cranial window implants in tree shrew and functional ultrasound imaging (fUSI) strategy. (A) Schematic of fUSI in tree shrew with visual stimulus presentation. (B-C) Schematic (B) and image (C) acoustic window and headplate on tree shrew. (D) Schematic of the chronic cranial window (coronal section) and of the ultrasound probe during imaging. Ultrasound gel is applied on top of the window, and the probe is positioned 2 mm above the window. (E) Schematic of the experimental setup. Hemodynamic signals are measured through the chronic cranial window using the ultrasound probe, stepped along the antero-posterior axis (black arrow). Three example image planes are labeled (numbered dashed lines). (F) Image of awake tree shrew in fUSI apparatus.

Compared to other imaging modalities commonly used in neuroscience, fUSI provides an attractive combination of depth penetration (several centimeters), high spatio-temporal resolution, and impressive sensitivity to hemodynamic fluctuations related to ongoing brain activity (Anwar et al., 2022; C. Rabut, Yoo, et al., 2020). While optical imaging offers the ability to visualize specific molecular signals such as intracellular calcium or voltage (Lee et al., 2016; Vanni, Thomas, Petry, Bickford, & Casanova, 2015; Wekselblatt, Flister, Piscopo, & Niell, 2016), it is limited in depth penetration due to the scattering properties of light. Meanwhile, functional magnetic resonance imaging (fMRI), provides noninvasive whole-brain access, but is limited by the requirement for MR-compatible non-metallic hardware and very high sensitivity to motion artifacts. Additionally, the non-portability and expense of the imaging magnet often limit availability of MR systems to researchers. Using fUSI, we image responses to classical visual stimuli, often used in fMRI to map visual cortical organization in other species. We find strong and spatially specific responses to retinotopy, dynamic noise, motion, object, and scrambled object stimuli in the tree shrew.

## Results

We developed chronic acoustic window implants that allow fUSI (Macé et al., 2011) from large volumes of the tree shrew cortex. This technique allowed us to image neural responses in successive coronal planes, the first corresponding to posterior V1 (just anterior of the transverse sinus), and approximately 6-7 mm of visibility lateral to the midline on each hemisphere. The data presented here have a lateral coverage of 12.8mm which is determined by the probe geometry, and more lateral areas are obscured by skull geometry. Successive planes were imaged relative to the lateral sinus as an anatomical landmark with various spacings. A schematic of our awake animals viewing various visual stimuli presented on a monitor positioned in front of the animal is shown in Figure 1A. We installed a large cranial window that enabled ultrasound waves to penetrate the brain for stable chronic imaging, as well as a headplate that allowed the animal to be held stationary without anesthesia (Figures 1B and 1C). We positioned the ultrasound probe above the cranial window and acquired a series of coronal images by stepping the probe along the antero-posterior axis, while the animal viewed different stimulus sets presented on a computer monitor placed in front of the eyes (Figures 1D, 1E and 1F). It should be noted that the imaging plane does not cover the whole lateral aspect of the tree shrew brain, and thus any extrastriate or inferotemporal visual areas lateral of ∼5mm of the midline would not be visible in the imaging plane without tilting the probe. Figures 4 and 5 correspond to the same animal in different imaging sessions, while figures 1, 2 and 3 come from different individual animals.

**Figure 2).**
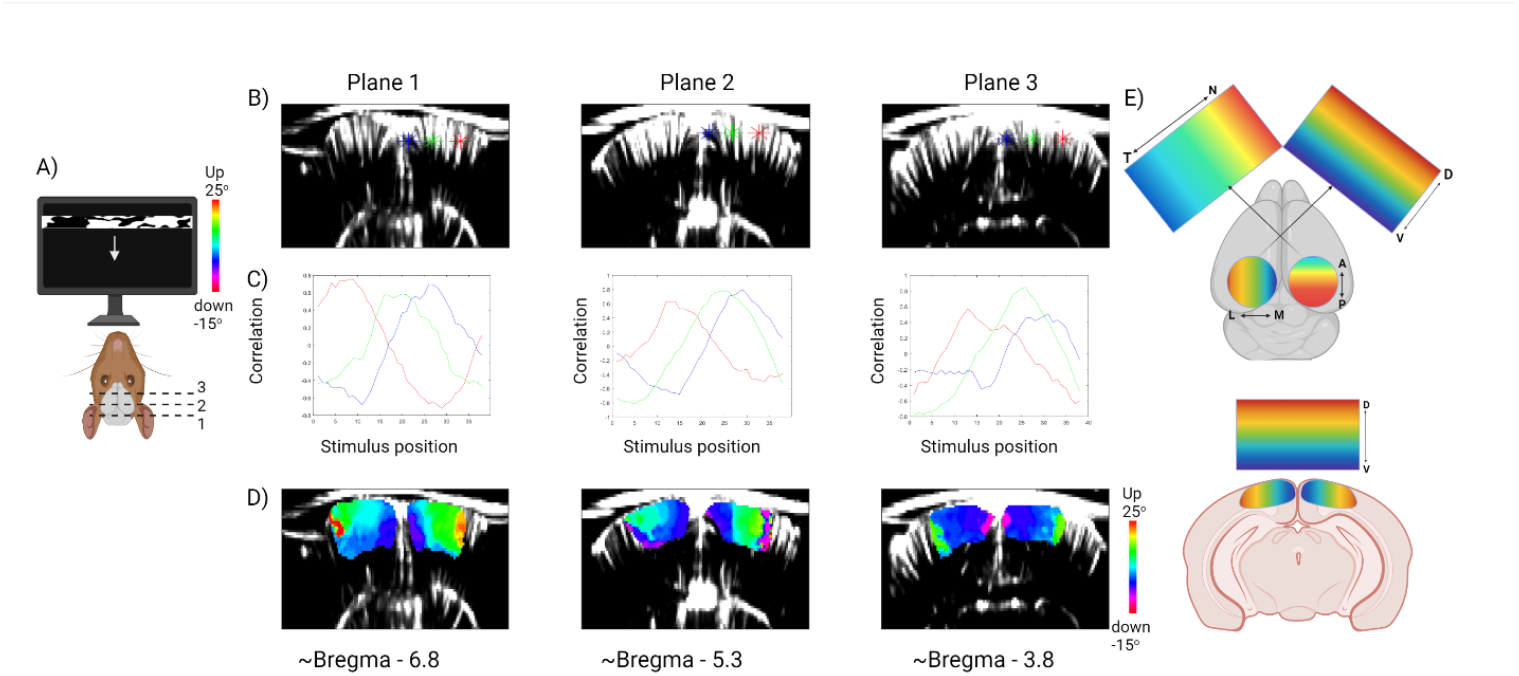
Retinotopic mapping in the tree shrew. (A) Schematic of the experiments for retinotopic mapping along the elevation axis. A horizontal bar is swept across the display monitors in front of the animal’s eyes (2°/s). Monitors were positioned so that each eye had a dedicated monitor meeting in front of the animal’s nose at a distance of 25 cm. Three example imaging planes crossing the primary visual cortex are labeled (numbered dashed lines). See also Figure S 2 and 3 for the azimuth retinotopic maps. (B) Example coronal slices showing the anatomical maps for the 3 planes shown by numbered dashed lines in panel A). Three voxels are marked in each plane (blue, green, and red stars) whose activity is shown by correlation with the retinotopic stimulus position in panel C). (C) Time-courses of correlation of the hemodynamic response as a function of stimulus position from the selected voxels shown in panel B). (D) Phase-maps in primary visual cortex for the three imaging planes, showing the preferred location of the horizontal bar on the screen shown in panel A). For this figure, all data is combined for trials with the bar moving up and trials with the bar moving down to compensate for hemodynamic delay. See Figure S 3 for the individual direction maps. E). Top – Schematic of observed retinotopic organization from the dorsal view with elevation represented on the left hemisphere / right screen and azimuth represented on the right hemisphere / left screen. Bottom – schematic of elevation retinotopic organization in V1 for the coronal view

**Figure 3).**
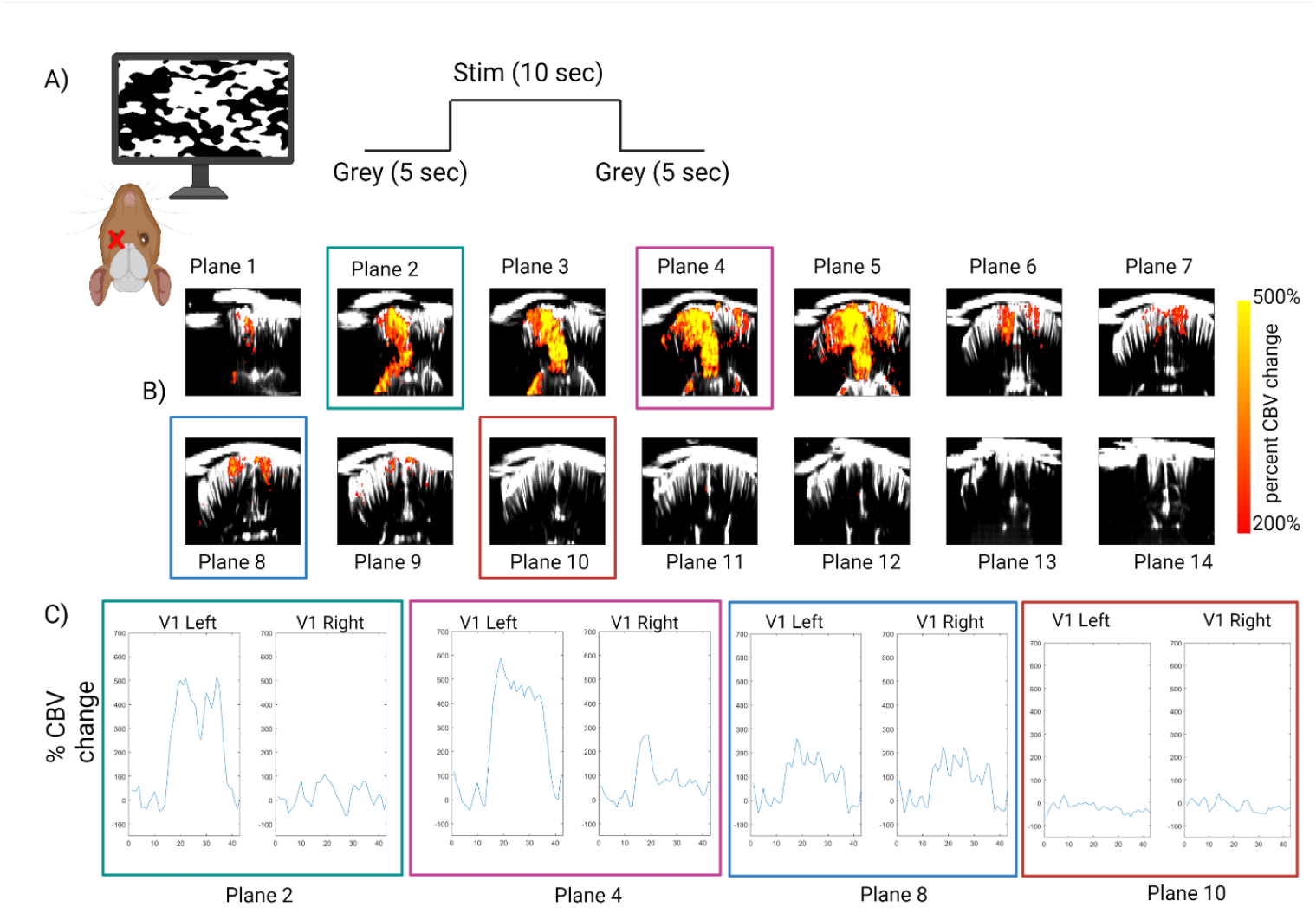
Response to full field noise stimulus using fUSI. (A) Schematic of the experiment for step function stimulus of 1/f binarized noise. Stimuli were presented to the right eye only and the timecourse of the stimulus function is shown on the right panel. (B) Activity maps overlaid on the anatomical image of the brain, thresholded at a 200% CBV change. Colored boxes indicate planes whose timecourses are shown in C). Planes are spaced by 0.75mm (C) Timecourse of activity for ROI’s of V1 on the left and right hemisphere for four example planes indicated in B).

**Figure 4).**
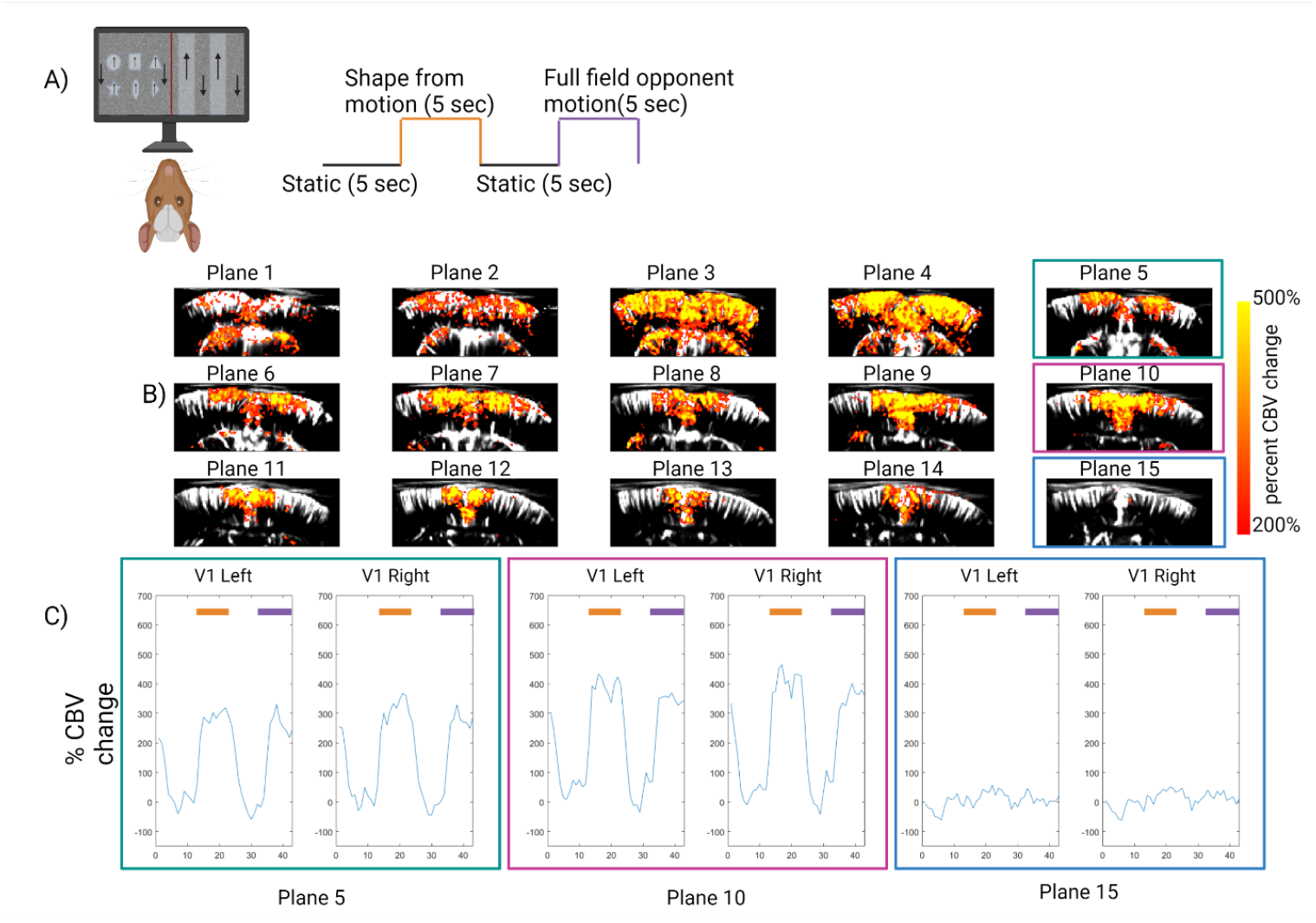
Response to motion stimulus using fUSI. (A) Schematic of the experiment for motion stimulus. Stimuli were presented to both eyes. The timecourse of the stimulus function is shown on the right panel. Two different kinds of motion were used, shape from motion (indicated in orange) and full field opponent motion (indicated in purple). (B) Activity maps overlaid on the anatomical image of the brain, thresholded at a 200% CBV change. Colored boxes indicate planes whose timecourses are shown in C). Planes are spaced by 0.4mm. (C) Timecourse of activity for ROI’s of V1 on the left and right hemisphere for three example planes indicated in B). Orange and purple bars indicate time of corresponding motion type (shape and opponent, respectively).

**Figure 5).**
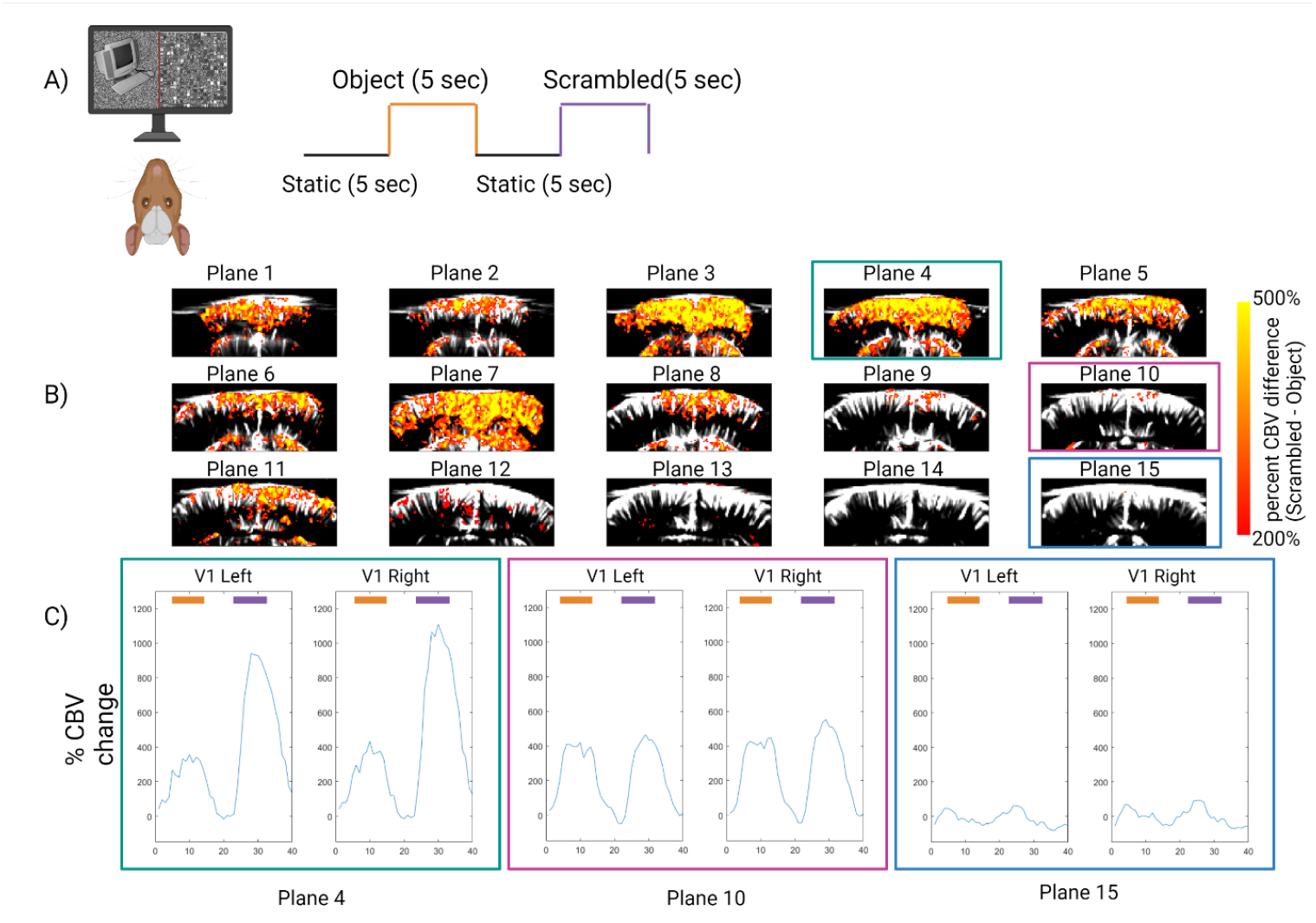
Response to object vs. scrambled stimulus using fUSI. (A) Schematic of the experiment for object vs. scrambled stimulus. Stimuli were presented to both eyes. The timecourse of the stimulus function is shown on the right panel. Object presentation is indicated in orange and scrambled object presentation is indicated in purple. (B) Activity difference maps (scrambled – object) are overlaid on the anatomical image of the brain, thresholded at a 200% CBV change. Colored boxes indicate planes whose timecourses are shown in C). (C) Timecourse of activity for ROI’s of V1 on the left and right hemisphere for three example planes indicated in B). Orange and purple bars indicate time of corresponding stimulus type (object and scrambled respectively).

### Retinotopic Mapping

Retinotopic maps are a primary organizing principle of primary visual cortex (V1) and other visually responsive areas such as the superior colliculus or extrastriate visual areas, where nearby points on the retina map to nearby points on the brain area, and orthogonal axes of the retinal surface are represented along orthogonal axes of the brain area. We first tested the ability of our ultrasound imaging system by performing retinotopic mapping with a stimulus consisting of a bar of binarized noise drifting either horizontally or vertically with a frequency *f* (Kalatsky and Stryker, 2003, Marshel, 2011). Here, we presented the stimulus to both eyes, each having a dedicated monitor positioned tangentially to the eye at 25cm distance, and we imaged responses from planes spaced at 1.5mm apart in the antero-posterior axis. Activity in visual cortex showed retinotopic organization (elevation, Figures 2A-2D and Supplemental Figures 3A and 3B; azimuth, Supplemental Figures 2, 3C and 3D). The activity induced by the retinotopic stimulus for different areas of primary visual cortex was highly correlated with different stimulus positions on the screen (Figure 2A-2D). This was especially evident in the elevation direction since changes in elevation maps were primarily seen in the medio-lateral axis captured by our successive coronal planes, spaced 1.5mm apart in the antero-posterior axis. We observed that the retinotopic map showed reversals in the elevation plane, corresponding to V1/V2 border (Supplemental figures 3A and 3B). These results indicate that fUSI provides a robust imaging method to perform retinotopic mapping experiments on individual animals, providing researchers with a tool to target recordings or manipulations to specific parts of the visual field. The stimuli here were not matched to the acuity of the animals and resolution of the maps can be improved in future work.. Here we restricted our analysis to cortical retinotopy but this can be extended to subcortical regions with retinotopy such as superior colliculus or the lateral geniculate nucleus of the thalamus. These coordinates can be used for fUS guided targeting of electrophysiology or viral injections to subfields of visual cortex or other functionally targeted regions of interest.

### Visual Responsiveness - Full Field Lava Stimulus

For mapping of total visual responsiveness, we then used a full field step function stimulus previously described by Niell and Stryker (Niell & Stryker, 2008), that contains 1/*f* binarized noise used for the retinotopy experiment above, but presented full field with interleaving grey periods. This is a high contrast stimulus containing edges of all orientations and moving in all directions to produce a large amount of visually evoked activity regardless of feature selectivity. This noise stimulus was presented to the right eye only on a dedicated monitor, positioned tangentially to the eye, in blocks of 10 seconds, with 5 seconds of gray screen before and after each cycle, for 10 cycles. We imaged 14 planes spaced 0.75 mm apart in the antero-posterior axis. This visual stimulus resulted in evoked activity in posterior areas, corresponding to the known location of visual cortex (Figure 3A-C). We also observed clear laterality in the response, which is especially strong in the more posterior sections, reflecting the contralateral coding of the right visual field by the left hemisphere visual cortex (Figure 3B-C). We computed the mean intensity during stimulus periods minus the mean during non-stimulus periods and then normalized to the mean during non-stimulus period to obtain the stimulus-evoked response in %CBV change (Figure 3B). We found values exceeding 5-fold changes in relative intensity with extremely high signal-to-noise, reflecting the value of the fUSI technique in identifying functional modules in the brain of the tree shrew. We also computed the correlation of the activity time course for each pixel with the stimulus time course, commonly used in functional ultrasound imaging analysis to reveal areas responsive to a given stimulus. We found that areas corresponding to the locations of superior colliculus (SC), primary visual cortex (V1) and secondary visual cortex (V2) all show high correlation values (>0.55), whereas non-visual areas are not correlated with the stimulus, ruling out effects of motion at stimulus onsets (Supplemental Figure 4).

### Motion Stimulus

As visual information proceeds along the visual pathway, the information that is represented and processed at each step is thought to become more complex. One such form of information is visual motion, and a variety of paradigms have been used to map the location of motion sensitive regions within the cortex of both humans and monkeys. The simplest form of a motion processing localizer involves the use of static versus moving dots. The contrast of moving dots versus static dots reveals a robust set of activations within the early visual system that is composed of areas MT, MST, and VIP in primates (Russ et al., 2021). For mapping of visual responses to motion, we used a standard block design format with a stimulus composed of two types of motion separated by static periods. Here, the screen was filled with low contrast dots on a grey background that either moved so that local opponent motion defined 6 shapes in the center portion of the screen, or which moved so that there was full field opponent motion of interleaving sections traveling up or down. This motion stimulus was presented to both eyes on a single monitor, positioned 25cm from the animal’s nose. Motion blocks were 5 seconds each, with 5 seconds of static before and after each presentation, for 10 cycles (Figure 4A). We imaged 15 planes spaced 0.4mm apart in the antero-posterior axis. This visual stimulus resulted in evoked activity in posterior areas (Figure 4B), corresponding to the known location of primary visual cortex, secondary visual area V2, and superior colliculus, with both types of motion evoking similar responses that were not significantly different in amplitude or activation pattern (Figure 4C). We did not observe any difference between the responses of the two hemispheres either. Since the responses to the different motion types did not differ significantly, we combined their responses for the activity maps seen in Figure 4B and the correlation map seen in Supplemental Figure 5. These results indicate that we can use fMRI-style localizers for mapping brain areas responsive to certain stimulus features, such as motion, using ultrasound imaging. This provides a much cheaper and more sensitive alternative to MRI experiments and can be easily adapted to other “stimulus sets and sensory systems. Unlike the specific motion responses seen in area MT of primates, in the tree shrew we observe that both primary visual cortex and extrastriate areas give strong motion responses despite no changes in luminance or contrast.

### Object vs. Scrambled Stimulus

Lastly, we compared the responses of objects to scrambled objects, a commonly used localizer in fMRI (Kourtzi & Kanwisher, 2000; Tsao, Freiwald, Knutsen, Mandeville, & Tootell, 2003). Here, we presented blocks of 10 objects on a noise background which changed at 2Hz for a period of 5 seconds, or grid scrambled versions of the same images, separated by 5 seconds of grey screen. This stimulus was presented to both eyes on a single monitor, positioned 25cm from the animal’s nose (Figure 5A). We imaged 15 planes spaced 0.4mm apart in the antero-posterior axis. This visual stimulus resulted in strong evoked activity for both stimuli (Supplemental Figures 6 and 7), but we found that the responses to the scrambled stimulus was generally significantly higher across the posterior portions of cortex, corresponding to primary visual cortex, and that no areas preferred the object stimulus to the scrambled (Figure 5B and C). This can likely be explained by the increased number of edges in the scrambled compared to object stimulus.

## Discussion

This study demonstrates the first brain-wide application of functional ultrasound imaging in the tree shrew and shows that fUSI enables high-resolution, high-sensitivity mapping of visual responses in awake animals. By combining chronic sonolucent cranial windows with coronal plane sampling, we produced reliable retinotopic maps, localized motion-sensitive regions, and contrasted object versus scrambled-object activations — all within single imaging sessions and with strong %CBV change signals across visual cortex and subcortical structures.

Two points of emphasis follow. First, fUSI fills a methodological gap between invasive single-unit electrophysiology and lower-resolution whole-brain modalities. Our maps provide the mesoscale context needed to (i) choose electrophysiological targets, (ii) interpret where local circuit recordings sit within a broader network, and (iii) deploy spatially precise perturbations (viral, chemogenetic, optical) across defined retinotopic loci. The combination of fUSI and targeted Neuropixels or optogenetic experiments will accelerate causal circuit dissection in this species.

Second, our results are complementary to recent Neuropixels work showing a compressed ventral visual hierarchy in tree shrew, with V2 exhibiting many properties typically associated with primate inferotemporal cortex. Whereas that electrophysiological study characterized receptive-field growth, latency changes and single-cell object representations, fUSI can potentially reveal where those computations are instantiated in the context of whole brain-level functional organization — including regions that are difficult to sample exhaustively with probes. Framing these datasets together provides a richer, multi-scale picture: high-density recordings reveal the microcircuit computations, and fUSI shows how those computations are distributed across the brain.

Practical advantages of the approach are notable: fUSI achieves fMRI-style whole-brain coverage with ∼100 µm resolution, is lower cost and more portable than MRI, and — with the cranial window approach — permits repeated chronic recordings in behaving animals. This makes fUSI particularly well suited to longitudinal studies (development, learning, plasticity), pharmacological manipulations, and combination with genetic or viral tools as those resources expand for tree shrew research.

Limitations and next steps: fUSI reports neurovascular signals, so complementary electrophysiology is essential to link hemodynamics to spiking dynamics and synaptic activity in any given region. Future studies should (i) perform simultaneous fUSI and Neuropixels/Calcium recordings to quantify the neurovascular transfer function across areas, (ii) use ethologically relevant stimulus sets and behavioral tasks to probe representations under naturalistic conditions, and (iii) apply targeted perturbations guided by fUSI maps to test causal roles of identified modules. Integrating mesoscale and single-cell datasets is the clearest path to answering whether compact stages like tree shrew V2 can, in fact, substitute for the broader primate IT in performing object recognition computations.

By establishing fUSI for brain-wide functional mapping in the tree shrew, we provide a practical, high-resolution method that links system-level maps to cellular-level studies. When combined with recent high-density electrophysiology and the growing genetic/viral toolkit, this approach positions the tree shrew as a uniquely useful model for studying visual computation at multiple scales — bridging rodent tractability and primate visual sophistication.

In conclusion, the northern tree shrew has shown potential as a model organism for the study of high-level vision and cognition. With its combination of high visual acuity, cone-dominant retina, and lissencephalic cortex, as well as its experimental and genetic accessibility, the tree shrew presents a unique opportunity to gain insight into the mechanisms of visual processing. The use of functional ultrasound imaging (fUSI) to record brain activity has allowed the mapping of functional modules of brain regions involved in visual perception, providing a valuable contribution to the field. This research opens up new avenues for further exploration of the visual processing pathways in the tree shrew and highlights its potential as a model for investigating other complex neural systems.

While the current study demonstrates the technical feasibility of high-resolution functional ultrasound imaging in the awake tree shrew, it is intended primarily as a methodological resource rather than a complete account of visual system organization. The results presented here were obtained from a limited number of animals and imaging sessions, and therefore should be viewed as proof-of-principle observations requiring replication and expansion. Future work incorporating alignment to a standard digital tree shrew atlas, quantitative flat-map analyses, and simultaneous or fUSI targeted electrophysiological recordings will be essential to establish the full reproducibility, coverage, and biological interpretation of the functional patterns observed here. The technical framework provided in this report is meant to enable such follow-up studies and to serve as a foundation for broader applications of fUSI in the tree shrew or other small-brain primate models such as marmosets.

## EXPERIMENTAL MODEL AND SUBJECT DETAILS

8 animals were used in this study, with 4 represented in the data presented here. Tree shrews (‘wild-type’, *Tupia belangari*) were bred onsite at Caltech. Original breeders were obtained from the Max Planc Florida Institute for Neuroscience and the Kunming Institute of Zoology. Animals were between six months and two years old. Both males and females were used. Animals were housed at 72-77 degrees F, 40-70% humidity, in a natural light cycle. Water and food pellets (ferret chow from Lab Diet) were provided ad libitum, with a daily rotation of fruits and vegetables. All animal procedures were performed in accordance with standard ethical guidelines (USDA) and were approved by the animal use committee (IACUC) at the California Institute of Technology.

## METHOD DETAILS

### Cranial Window Implantation

Tree shrews were anesthetized with fentanyl-midazolam-dexdomitor (FMD, fentanyl 0.05 mg/kg, midazolam 5.0 mg/kg, dexdomitor 0.25 mg/kg, injected subcutaneously (s.c.). Scalp hair was then removed using an electric shaver and hair removal lotion (Nair) was used to expose the skin for the surgical procedure. Topical Lidocaine hydrochloride jelly (2%; Akorn, Lake Forest, IL) was applied to the top of the scalp and ears and left for 1–2 min before being cleared with isopropanol. Chlorhexadine solution was then applied to the surgical site with cotton-tipped applicators for final sterilization. Bupivicaine 0.5% was injected just under the scalp and left for at least 1 minute before any incisions. Animals were positioned in a stereotax using ear bars placed just below the ear canal for stability. Eye gel (Puralube) was applied to prevent corneal dehydration during surgery. The animals were given subcutaneous injections of the analgesic Ketoprofen (5 mg/kg), an antibiotic Baytril (5 mg/kg), a steroidal anti-inflammatory Dexamethasone (5mg/kg), an osmotic diuretic Mannitol 20% (.8ml) to prevent brain swelling, and 2 ml saline to prevent postoperative dehydration. Body temperature was maintained at 38.5°C by a feedback-controlled heating pad; temperature, heart rate, SpO2, and breathing were monitored throughout surgery and recorded at least every 15 minutes. Sterilized instruments and aseptic technique was used throughout.

Following a scalp incision, the periosteum was cleared from the surface of the skull using #3 forceps. Once the skull was completely cleared of connective tissue, the edges of the skin incision were attached to the skull with tissue adhesive (Vetbond, veterinary grade, 3M) leaving room for the subsequent craniotomy and headplate. A large craniotomy was made with a drill (0.5 to 0.7 mm tip) extending from bregma to lambda along the antero-posterior axis, and from -7 to +7 mm from the midline along the lateral axis. To prevent heating of cortex, the drilling of the craniotomy was done slowly, using compressed air to blow off bone chips with a cold, sterile saline rinse between short periods of drilling. When drilling across the sutures, care was taken to avoid damaging the underlying sinuses. Once the inner portion of the craniotomy moved somewhat freely when touched, angled forceps were used to remove the center piece of bone carefully, lifting vertically while the skull was immersed in saline. Dura was kept intact. After removal of the skull, a thin layer of transparent polymer (part no. 3-4680; Dow Corning, Midland, MI), used to seal craniotomies in primates (Jackson & Muthuswamy, 2008), was applied to the dura surface, just enough to cover the craniotomy, for protection and stability. Next, a flap of sonolucent plastic (Polymethylpentene, 250 mm thick, Goodfellow) was cut to fit the cranial window, positioned on top of a layer of the artificial dura, and sealed with tissue adhesive (Supplemental Figure 1).

A custom titanium head bar was attached to the front of the skull with tissue adhesive (Vetbond, veterinary grade, 3M) and dental cement (C&B Metabond) with care taken to fully seal the headplate to the skull on all sides while avoiding getting adhesive on the window. Anesthesia was reversed with atipamezole-flumazenil (s.c., atipamezole 1.25 mg/kg, flumazenil 0.25 mg/kg). Animals were left to recover for at least 5 days before the first imaging session. The surgery was adapted from published protocols of acoustic windows in mice (Brunner et al., 2020; Mace et al., 2018) and from protocols of large chronic cranial window implantations used for optical imaging with glass (Wekselblatt et al., 2016).

### Functional Ultrasound Imaging

For each individual experimental session, an awake tree shrew with a cranial window was head-fixed in front of two stimulation monitors and the body was restrained in a 3D printed tube. Acoustic coupling gel (1 mL) was applied on the cranial window for ultrasound coupling before placing the ultrasound probe. The acoustic gel was centrifuged to avoid air bubbles, and warmed to body temperature in a water bath before application to the window. The ultrasound probe (Domino, Vermon) was positioned into the gel ∼2 mm above the cranial window (Figure 1F). Gel was re-applied about every hour without removing the ultrasound probe. The probe holder was mounted on a linear micro precision motor (Zaber) that moved the probe holder along the antero-posterior axis. The probe was connected to an ultrasound scanner (Vantage 128, Verasonics) controlled by a PC. Visual stimuli were presented while functional ultrasound images were taken (2 Hz). Visual stimulation was presented using Matlab (psychtoolbox) with a frame rate of 30Hz. For experiments using a single monitor, stimuli were presented with a screen (27’’ ViewSonic VA2759-SMH 27) positioned directly in front of the animal at 0° with regard to the nose. For retinotopy experiments, there were two monitors placed on each side of the animal’s head in landscape orientation tangentially to each eye, at an angle of 45° with regard to the antero-posterior axis of the tree shrew. The monitors were placed 25 cm away from the animal.

### Visual stimuli

A visual stimulation block was composed of alternating gray backgrounds (10 s each, before and after visual stimuli), and either retinotopy (in one of four directions) 1/*f* full-field binarized noise, motion or object stimuli. different directions (right, up, left or down). The stimulation block was repeated 10 times at each coronal plane.

### Analysis of Visual Responsiveness: Motion, Full Field Lava and Object vs Shape stimuli

To examine the activation patterns evoked by the three non-retinotopic stimuli used in this paper, both correlation and activation maps were estimated. These maps provided a spatial representation of brain regions that exhibited significant changes in activity during the presentation of each stimulus.

#### Data Pre-processing

Prior to the analysis, the fUSI data underwent preprocessing steps to correct for drifts in the hemodynamic baseline and obtain a representative dataset. The recorded fUSI data was processed using the MATLAB function ‘smooth’ with a large kernel size of 25% of the temporal dimension. This correction method specifically targeted long-term, slow-changing temporal fluctuations while preserving the stimulus response. The data from all ten iterations were then averaged to create a single representative dataset, enhancing signal quality and reducing noise and artifacts. This averaged dataset served as the foundation for further analysis, enabling a focused examination of the correlation between the stimuli and fUSI signals.

#### Estimation of Correlation Map

To evaluate the relationship between each stimulus and the fUSI signals, a cross-correlation (CC) analysis was conducted. This analysis aimed to measure the similarity or correlation between the stimulus and the fUSI signals at each spatial pixel in the dataset. By applying the cross-correlation function to the data, cross-correlation maps were generated. For the cross-correlation analysis, the on-off activation function was aligned with the stimulus events, considering a time shift of 3 frames. This time shift factor accounted for the rise-time of the response and ensured accurate alignment for effective correlation analysis. Multiple correlation maps were generated, capturing different combinations of the stimulus conditions. Each map provided valuable information for further analysis and investigation of the relationship between the stimuli and the fUSI signals.

#### Estimation of Activation Map

To investigate the activation patterns evoked by each stimulus, activation maps were estimated. These maps provided a spatial representation of brain regions that exhibited significant changes in activity during the presentation of the stimulus. To calculate the activation map, specific time frames corresponding to the ON and OFF periods of each stimulus were identified. The first step involved subtracting the minimum value of each pixel across the entire timeline, normalizing the data to focus on relative changes in activity. The activation maps were generated by computing the mean signal values during the ON and OFF periods. For each stimulus condition, the activation map was obtained by calculating the percentage change between the mean signal during the ON period (mean of all ON frames) and the mean signal during the OFF period (mean of all OFF frames). This percentage change was then multiplied by 100 and divided by the mean signal during the OFF period.

By performing these steps for each stimulus, the paper was able to examine the correlation between the stimuli and fUSI signals, identify brain regions that responded to each stimulus, and compare the responses during the ON and OFF periods. This comprehensive analysis provided valuable insights into the neural activity patterns evoked by each of the four different stimuli.

### Retinotopic Mapping

The analysis of the retinotopy data involved several steps. Initially, a reference baseline frame was calculated by averaging the 300 fUSI frames acquired before the visual stimulus. Spatial median filtering was then applied in the axial and lateral directions with kernel sizes of 3 and 4 pixels, respectively, to improve frame quality and reduce noise. Subsequently, the fUSI frames were averaged across 20 repetitions of the stimulus function, combining opposing directions such as left-to-right with reversed right-to-left and top-to-bottom with bottom-to-top retinotopic stimulation. This averaging step was employed to enhance the signal-to-noise ratio, promoting a more accurate and reliable analysis of the retinotopy data.

To analyze the correlation and estimate the phase maps, a stimulation function representing the ON and OFF periods of the stimulus was created and circularly shifted to capture different temporal phases. Correlation maps were generated by measuring the temporal similarity between the applied stimulation and the stimulus-induced activity. The mean correlation motion was subtracted from the entire movie to enhance activation contrast.

For phase map generation, the indices corresponding to the maximum values at each pixel in the correlation dataset were determined. These indices represented the temporal phases evoking the strongest responses in the fUSI data. To improve clarity and reduce noise, a 2×2 median filtering was applied to the phase maps. This process emphasized the underlying retinotopic patterns and facilitated data interpretation. Phase maps were generated consistently for both vertical and horizontal visual stimulus conditions, enabling a comprehensive analysis of the retinotopy data and cortical organization exploration.

## Acknowledgements

We are deeply grateful to Doris Y. Tsao and Mikhail G. Shapiro for their mentorship and financial support of this and related tree shrew projects exploring a range of neurotechnologies in this valuable research model. We thank Daniel A. Wagenaar and the Caltech Neurotechnology Laboratory for contributions to the design and manufacturing of the head-fixation system. We thank Rafael Grytz at the University of Alabama at Birmingham for world-class support on tree shrew husbandry practices. We thank Nicole Schweers, Audo Flores, and Margaret Swift for expert lab management.

## Author Contributions

Conceptualization was performed by J.B.W., F.F.L. Software for visual stimulus presentation and synchronization was developed by F.F.L. and F.L. Formal analysis was conducted by R.N. Experiments were performed by J.B.W., who also operated the tree shrew colony. Surgeries were carried out by J.B.W and F.F.L.. The original draft was written by J.B.W. and R.N.

## References

Anwar, O. N.-E., Michael, K., Charu Bai, R., Gabriel, M., Alan, U., Kenneth, D. H., & Matteo, C. (2022). Neural correlates of blood flow measured by ultrasound. Neuron, 110(10), 1631-1640.e1634. doi:10.1016/j.neuron.2022.02.012

Bimbard, C., Demene, C., Girard, C., Radtke-Schuller, S., Shamma, S., Tanter, M., & Boubenec, Y. (2018). Multi-scale mapping along the auditory hierarchy using high-resolution functional UltraSound in the awake ferret. Elife, 7. doi:10.7554/eLife.35028

Brunner, C., Grillet, M., Sans-Dublanc, A., Farrow, K., Lambert, T., Mace, E., … Urban, A. (2020). A Platform for Brain-wide Volumetric Functional Ultrasound Imaging and Analysis of Circuit Dynamics in Awake Mice. Neuron, 108(5), 861–875 e867. doi:10.1016/j.neuron.2020.09.020

Chan, K. Y., Jang, M. J., Yoo, B. B., Greenbaum, A., Ravi, N., Wu, W. L., … Gradinaru, V. (2017). Engineered AAVs for efficient noninvasive gene delivery to the central and peripheral nervous systems. Nat Neurosci, 20(8), 1172–1179. doi:10.1038/nn.4593

Chuapoco, M. R., Flytzanis, N. C., Goeden, N., Octeau, J. C., Roxas, K. M., Chan, K. Y., … Gradinaru, V. (2022). Intravenous gene transfer throughout the brain of infant Old World primates using AAV. bioRxiv, 2022.2001.2008.475342. doi:10.1101/2022.01.08.475342

Demene, C., Baranger, J., Bernal, M., Delanoe, C., Auvin, S., Biran, V., … Baud, O. (2017). Functional ultrasound imaging of brain activity in human newborns. Sci Transl Med, 9(411). doi:10.1126/scitranslmed.aah6756

Demené, C., Bimbard, C., Gesnik, M., Radtke-Schuller, S., Shamma, S., Boubenec, Y., & Tanter, M. (2016, 18-21 Sept. 2016). Functional Ultrasound Imaging of the thalamo-cortical auditory tract in awake ferrets using ultrafast Doppler imaging. Paper presented at the 2016 IEEE International Ultrasonics Symposium (IUS).

Gawne, T. J., Siegwart, J. T., Jr., Ward, A. H., & Norton, T. T. (2017). The wavelength composition and temporal modulation of ambient lighting strongly affect refractive development in young tree shrews. Experimental eye research, 155, 75–84. doi:10.1016/j.exer.2016.12.004

Gawne, T. J., Ward, A. H., & Norton, T. T. (2018). Juvenile Tree Shrews Do Not Maintain Emmetropia in Narrow-band Blue Light. Optom Vis Sci, 95(10), 911–920. doi:10.1097/opx.0000000000001283

Heidenreich, M., & Zhang, F. (2016). Applications of CRISPR-Cas systems in neuroscience. Nat Rev Neurosci, 17(1), 36–44. doi:10.1038/nrn.2015.2

Hu, W., Zhu, S., Briggs, F., & Doyley, M. M. (2023). Functional ultrasound imaging reveals 3D structure of orientation domains in ferret primary visual cortex. Neuroimage, 268, 119889. doi:10.1016/j.neuroimage.2023.119889

Imbault, M., Chauvet, D., Gennisson, J. L., Capelle, L., & Tanter, M. (2017). Intraoperative Functional Ultrasound Imaging of Human Brain Activity. Sci Rep, 7(1), 7304. doi:10.1038/s41598-017-06474-8

Jackson, N., & Muthuswamy, J. (2008). Artificial dural sealant that allows multiple penetrations of implantable brain probes. J Neurosci Methods, 171(1), 147–152. doi:10.1016/j.jneumeth.2008.02.018

Jang, J., Song, M., & Paik, S.-B. (2019). Classification of columnar and salt-and-pepper organization in mammalian visual cortex. bioRxiv, 698043. doi:10.1101/698043

Kourtzi, Z., & Kanwisher, N. (2000). Cortical regions involved in perceiving object shape. J Neurosci, 20(9), 3310–3318. doi:10.1523/JNEUROSCI.20-09-03310.2000

Lander, E. S. (2016). The Heroes of CRISPR. Cell, 164(1-2), 18-28. doi:10.1016/j.cell.2015.12.041

Lanfranchi, F., Wekselblatt, J., Wagenaar, D., Tsao, D. (2025). A compressed hierarchy for visual form processing in the tree shrew. Nature (2025). 10.1038/s41586-025-09441-w

Lee, K. S., Huang, X., & Fitzpatrick, D. (2016). Topology of ON and OFF inputs in visual cortex enables an invariant columnar architecture. Nature, 533(7601), 90–94. doi:10.1038/nature17941

Lyon, D. C., Jain, N., & Kaas, J. H. (1998). Cortical connections of striate and extrastriate visual areas in tree shrews. J Comp Neurol, 401(1), 109–128. doi:10.1002/(sici)1096-9861(19981109)401:1<109::aid-cne7>3.0.co;2-i

Macé, E., Montaldo, G., Cohen, I., Baulac, M., Fink, M., & Tanter, M. (2011). Functional ultrasound imaging of the brain. Nature Methods, 8(8), 662–664. doi:10.1038/nmeth.1641

Mace, E., Montaldo, G., Trenholm, S., Cowan, C., Brignall, A., Urban, A., & Roska, B. (2018). Whole-Brain Functional Ultrasound Imaging Reveals Brain Modules for Visuomotor Integration. Neuron, 100(5), 1241–1251 e1247. doi:10.1016/j.neuron.2018.11.031

Madineh, S.-S., Kuo-Sheng, L., Juliane, J., Rachel, S., Nicole, S., & David, F. (2021). A sinusoidal transformation of the visual field is the basis for periodic maps in area V2. Neuron, 109(24), 4068-4079.e4066. doi:10.1016/j.neuron.2021.09.053

Niell, C. M., & Stryker, M. P. (2008). Highly selective receptive fields in mouse visual cortex. J Neurosci, 28(30), 7520–7536. doi:10.1523/jneurosci.0623-08.2008

Norman, S. L., Maresca, D., Christopoulos, V. N., Griggs, W. S., Demene, C., Tanter, M., … Andersen, R. A. (2020). Single Trial Decoding of Movement Intentions Using Functional Ultrasound Neuroimaging. bioRxiv, 2020.2005.2012.086132. doi:10.1101/2020.05.12.086132

Petry, H. M., & Bickford, M. E. (2019). The Second Visual System of The Tree Shrew. J Comp Neurol, 527(3), 679–693. doi:10.1002/cne.24413

Petry, H. M., Fox, R., & Casagrande, V. A. (1984). Spatial contrast sensitivity of the tree shrew. Vision Res, 24(9), 1037–1042. doi:10.1016/0042-6989(84)90080-4

Rabut, C., Correia, M., Finel, V., Pezet, S., Pernot, M., Deffieux, T., & Tanter, M. (2019). 4D functional ultrasound imaging of whole-brain activity in rodents. Nature Methods, 16(10), 994–997. doi:10.1038/s41592-019-0572-y

Rabut, C., Ferrier, J., Bertolo, A., Osmanski, B., Mousset, X., Pezet, S., … Tanter, M. (2020). Pharmaco-fUS: Quantification of pharmacologically-induced dynamic changes in brain perfusion and connectivity by functional ultrasound imaging in awake mice. Neuroimage, 222, 117231. doi:10.1016/j.neuroimage.2020.117231

Rabut, C., Yoo, S., Hurt, R. C., Jin, Z., Li, H., Guo, H., … Shapiro, M. G. (2020). Ultrasound Technologies for Imaging and Modulating Neural Activity. Neuron, 108(1), 93–110. doi:10.1016/j.neuron.2020.09.003

Russ, B. E., Petkov, C. I., Kwok, S. C., Zhu, Q., Belin, P., Vanduffel, W., & Hamed, S. B. (2021). Common functional localizers to enhance NHP & cross-species neuroscience imaging research. Neuroimage, 237, 118203. doi:10.1016/j.neuroimage.2021.118203

Sajdak, B. S., Salmon, A. E., Cava, J. A., Allen, K. P., Freling, S., Ramamirtham, R., Carroll, J. (2019). Noninvasive imaging of the tree shrew eye: Wavefront analysis and retinal imaging with correlative histology. Exp Eye Res, 185, 107683. doi:10.1016/j.exer.2019.05.023

Schumacher, J. W., McCann, M., Maximov, K. J., & Fitzpatrick, D. (2021). Selective enhancement of neural coding in V1 underlies fine discrimination learning in tree shrew. bioRxiv, 2021.2001.2010.426145. doi:10.1101/2021.01.10.426145

Sedigh-Sarvestani, M., & Fitzpatrick, D. (2022). What and Where: Location-Dependent Feature Sensitivity as a Canonical Organizing Principle of the Visual System. Frontiers in Neural Circuits, 16. doi:10.3389/fncir.2022.834876

Sieu, L. A., Bergel, A., Tiran, E., Deffieux, T., Pernot, M., Gennisson, J. L., Cohen, I. (2015). EEG and functional ultrasound imaging in mobile rats. Nat Methods, 12(9), 831–834. doi:10.1038/nmeth.3506

Sternberg, S. H., & Doudna, J. A. (2015). Expanding the Biologist’s Toolkit with CRISPR-Cas9. Mol Cell, 58(4), 568–574. doi:10.1016/j.molcel.2015.02.032

Tiran, E., Ferrier, J., Deffieux, T., Gennisson, J. L., Pezet, S., Lenkei, Z., & Tanter, M. (2017). Transcranial Functional Ultrasound Imaging in Freely Moving Awake Mice and Anesthetized Young Rats without Contrast Agent. Ultrasound Med Biol, 43(8), 1679–1689. doi:10.1016/j.ultrasmedbio.2017.03.011

Topcuoglu, M. A., Aydin, H., & Saka, E. (2009). Occipital cortex activation studied with simultaneous recordings of functional transcranial Doppler ultrasound (fTCD) and visual evoked potential (VEP) in cognitively normal human subjects: effect of healthy aging. Neurosci Lett, 452(1), 17–22. doi:10.1016/j.neulet.2009.01.030

Tsao, D. Y., Freiwald, W. A., Knutsen, T. A., Mandeville, J. B., & Tootell, R. B. (2003). Faces and objects in macaque cerebral cortex. Nat Neurosci, 6(9), 989–995. doi:10.1038/nn1111

Urban, A., Mace, E., Brunner, C., Heidmann, M., Rossier, J., & Montaldo, G. (2014). Chronic assessment of cerebral hemodynamics during rat forepaw electrical stimulation using functional ultrasound imaging. Neuroimage, 101, 138–149. doi:10.1016/j.neuroimage.2014.06.063

Vanni, M. P., Thomas, S., Petry, H. M., Bickford, M. E., & Casanova, C. (2015). Spatiotemporal Profile of Voltage-Sensitive Dye Responses in the Visual Cortex of Tree Shrews Evoked by Electric Microstimulation of the Dorsal Lateral Geniculate and Pulvinar Nuclei. J Neurosci, 35(34), 11891–11896. doi:10.1523/JNEUROSCI.0717-15.2015

Ward, A. H., Norton, T. T., Huisingh, C. E., & Gawne, T. J. (2018). The hyperopic effect of narrow-band long-wavelength light in tree shrews increases non-linearly with duration. Vision Research, 146-147, 9–17. doi:10.1016/j.visres.2018.03.006

Wekselblatt, J. B., Flister, E. D., Piscopo, D. M., & Niell, C. M. (2016). Large-scale imaging of cortical dynamics during sensory perception and behavior. J Neurophysiol, 115(6), 2852–2866. doi:10.1152/jn.01056.2015

Wong, P., & Kaas, J. H. (2009). Architectonic subdivisions of neocortex in the tree shrew (Tupaia belangeri). Anat Rec (Hoboken), 292(7), 994–1027. doi:10.1002/ar.20916

Xiao, J., Liu, R., & Chen, C. S. (2017). Tree shrew (Tupaia belangeri) as a novel laboratory disease animal model. Zool Res, 38(3), 127–137. doi:10.24272/j.issn.2095-8137.2017.033

