## supplemental figures for "Mapping function in the tree shrew visual system using functional ultrasound imaging"

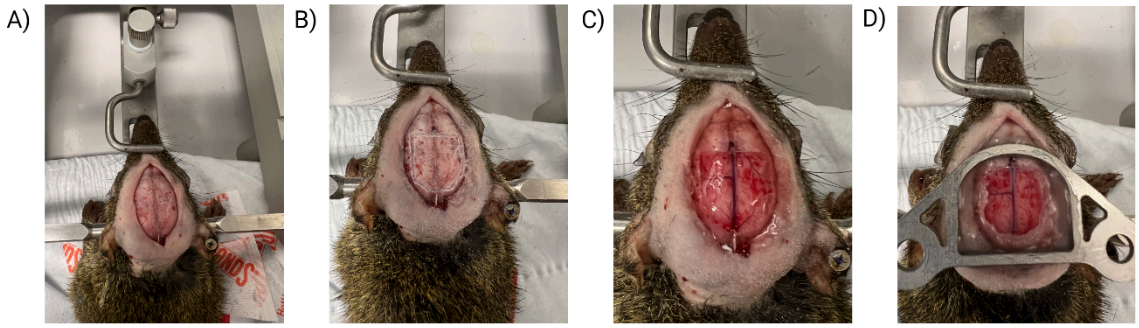


**Supplemental Figure 1). Surgical preparation. (**A) Animal is positioned on stereotaxic frame. After incision of ~1cm, skull is leveled and connective tissue should be cleared away. (B) The position of Bregma is marked and the outline of the window implant is drilled slowly so not to create excess heating. (C) The skull is removed carefully and a TPX window of the same shape is placed on the brain (0.125 mm thickness Polymethylpentene). An optional layer of artificial dura may be placed on the cortical surface to protect the brain from adhesive and help prevent regrowth (3−4680, Dow Corning, Midland, MI). (D) A headplate is sealed to the skull using cyanoacrylate and dental cement (C&B metabond).

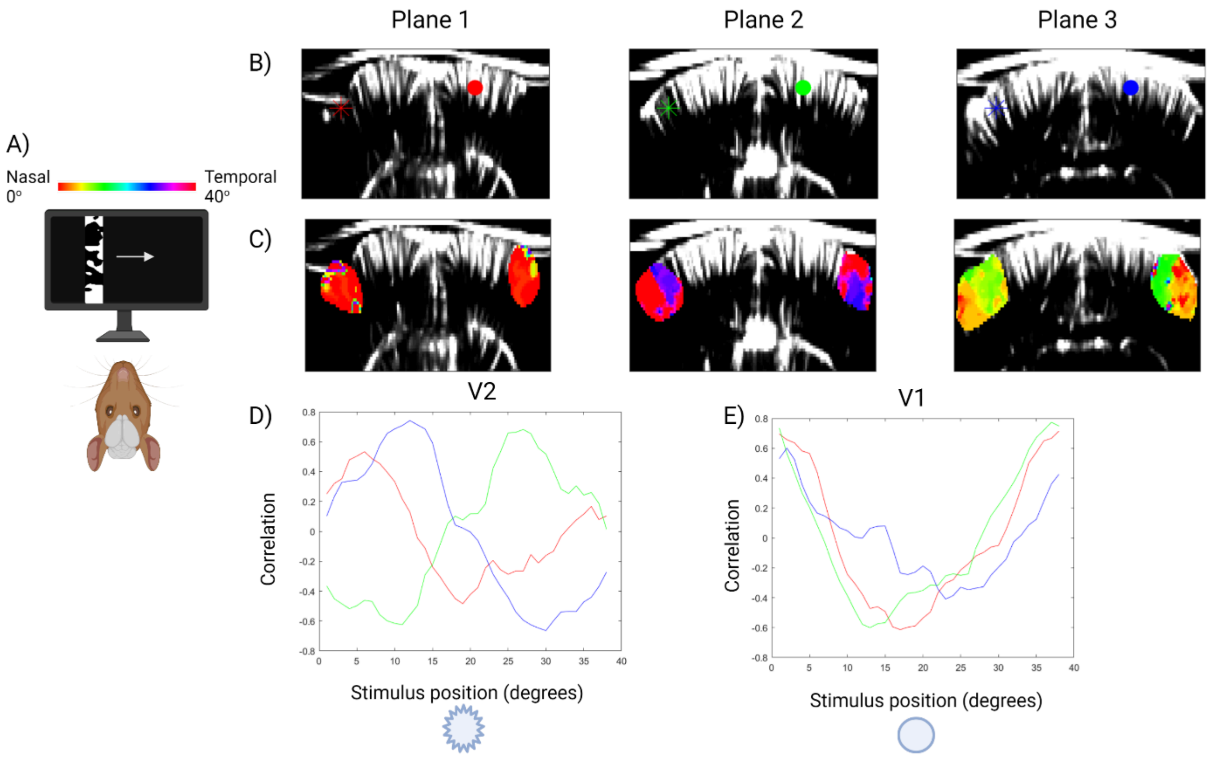


**Supplemental Figure 2). Retinotopic mapping of azimuth in the tree shrew.** (A) Schematic of the experiments for retinotopic mapping along the azimuth axis. A vertical bar is swept across the display monitors in front of the animals' eyes (2°/s). Monitors were positioned so that each eye had a dedicated monitor meeting in front of the animal's nose at 25cm.distance. (B) Example coronal slices showing the anatomical maps for 3 planes spaced by 1.5mm (posterior to anterior). One voxel is marked in V2 each plane (stars colored by plane) whose activity is shown by correlation with the retinotopic stimulus position in panel D), and another voxel is marked in V1 each plane (circles colored by plane) whose activity is shown by correlation with the retinotopic stimulus position in panel E). (D) Phase-maps in secondary visual area (V2) for the three imaging planes, showing the preferred location of the horizontal bar on the screen shown in panel A). (D) Time-courses of correlation of the hemodynamic response as a function of stimulus position from the selected voxels for V2 shown by stars in panel B). (E) Time-courses of correlation of the hemodynamic response as a function of stimulus position from the selected voxels for V1 shown by circles in panel B).

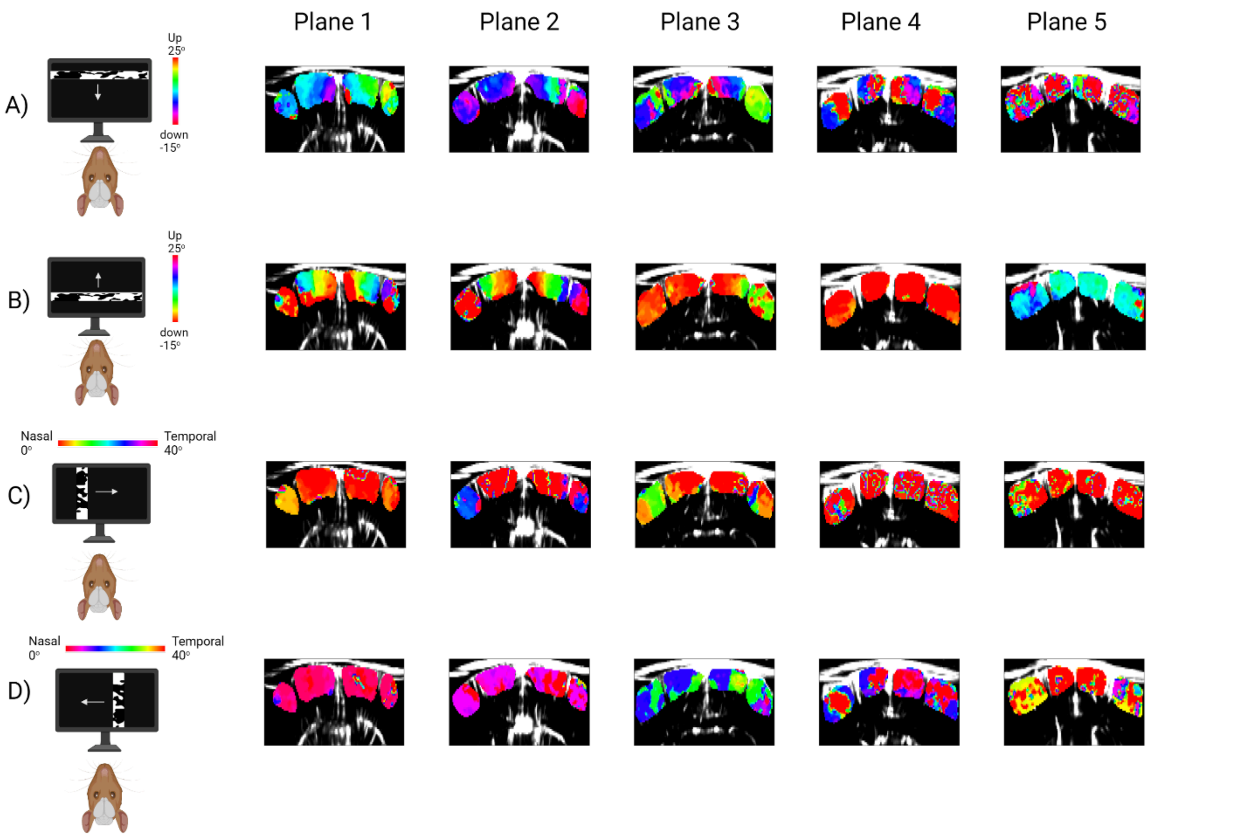


**Supplemental Figure 3). Retinotopic maps by direction.** (A) Elevation phase-maps in primary visual cortex (V1) and secondary visual area (V2) for the five imaging planes, showing the preferred location of the horizontal bar on the screen moving from top-to-bottom. (B) Elevation phase-maps as in A), but with the bar moving from bottom-to-top. (C) Azimuth phase-maps in primary visual cortex (V1) and secondary visual area (V2) for the five imaging planes, showing the preferred location of the vertical bar on the screen moving from nasal-to-temporal. (D) Azimuth phase-maps as in C), but with the bar moving from temporal-to-nasal.

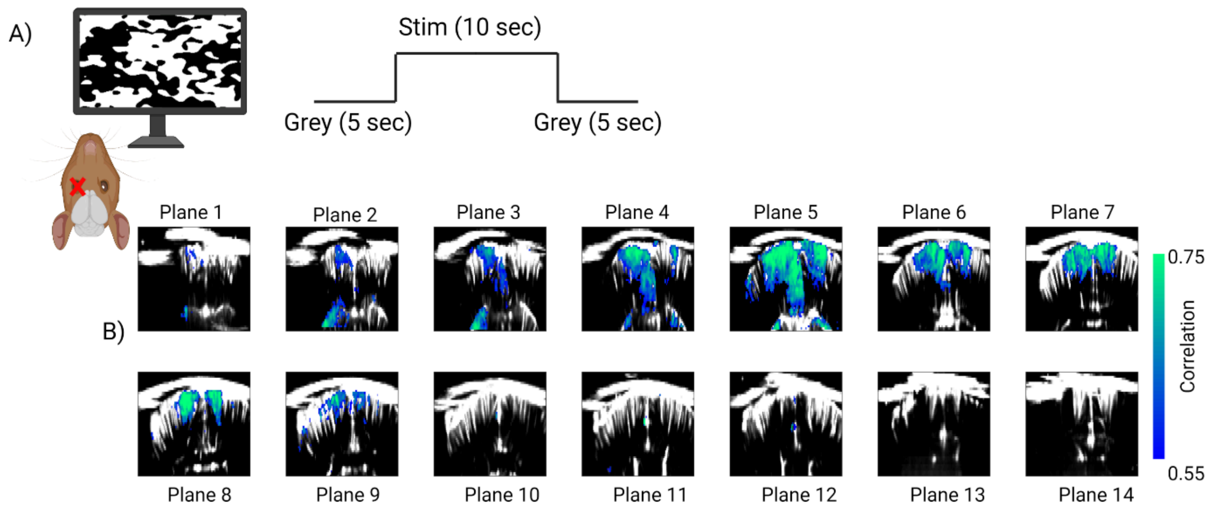


**Supplemental Figure 4). Correlation map to full field noise stimulus** (A) Schematic of the experiment for step function stimulus of 1/f binarized noise. Stimuli were presented to the right eye only and the timecourse of the stimulus function is shown on the right panel. (B) Correlation maps overlaid on the anatomical image of the brain, thresholded at a correlation value of 0.55. Planes are spaced by 0.75mm.

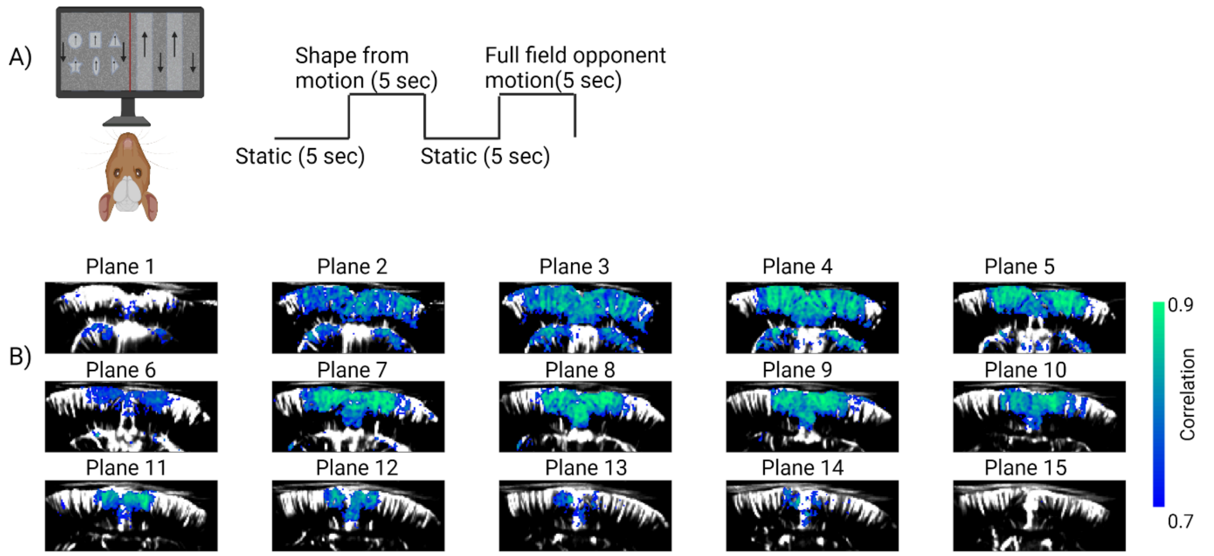


**Supplemental Figure 5). Correlation map to motion stimulus** (A) Schematic of the experiment for motion stimulus. Stimuli were presented to the both eyes and the timecourse of the stimulus function is shown on the right panel. (B) Correlation maps overlaid on the anatomical image of the brain, thresholded at a correlation value of 0.7. Planes are spaced by 0.4mm

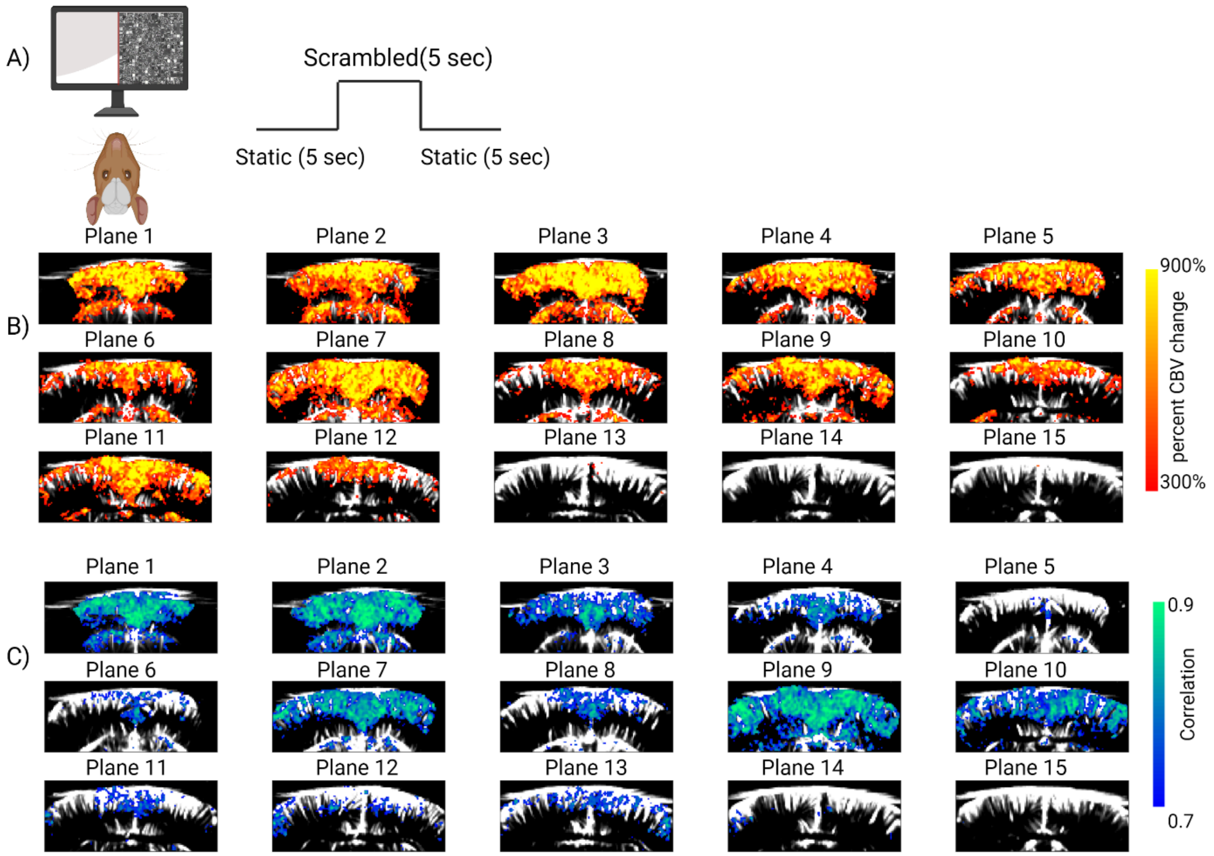


**Supplemental Figure 6). Response to scrambled objects using fUSI** (A) Schematic of the experiment for object stimulus. Stimuli were presented to the both eyes and the timecourse of the stimulus function is shown on the right panel. (B) Activity maps overlaid on the anatomical image of the brain, thresholded at a 300% CBV change. Planes are spaced by 0.4mm. (C) Correlation maps overlaid on the anatomical image of the brain, thresholded at a correlation value of 0.7.

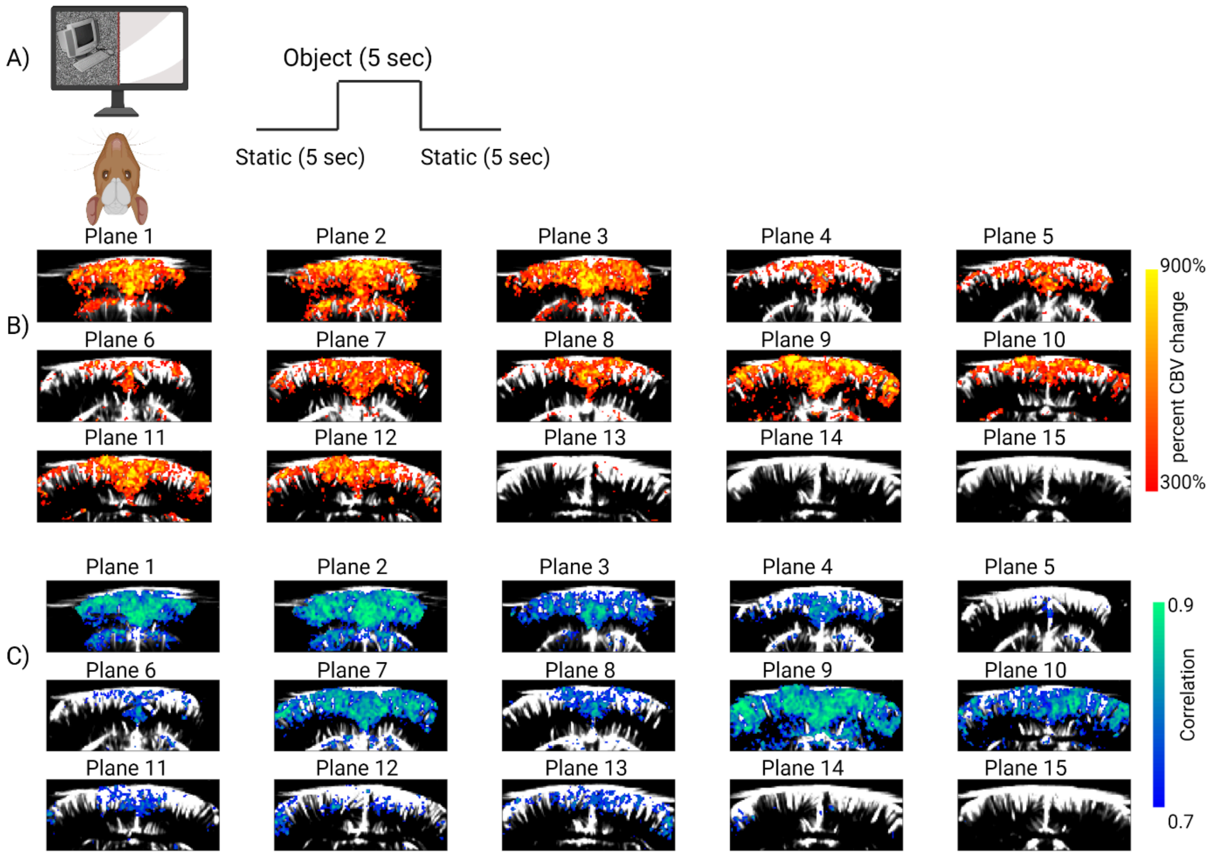


**Supplemental Figure 7). Response to objects using fUSI** (A) Schematic of the experiment for scrambled stimulus. Stimuli were presented to the both eyes and the timecourse of the stimulus function is shown on the right panel. (B) Activity maps overlaid on the anatomical image of the brain, thresholded at a 300% CBV change. Planes are spaced by 0.4mm. (C) Correlation maps overlaid on the anatomical image of the brain, thresholded at a correlation value of 0.7.
